# Intermittent Hypoxia Alters Cerebrovascular Recovery After Stroke

**DOI:** 10.64898/2026.01.11.698922

**Authors:** Bayan El Amine, Aurélien Delphin, Marc-Adrien Reveyaz, Célian Peyronnel, Emeline Lemarié, Nora Collomb, Sophie Bouyon, Antoine Boutin-Paradis, Hisham Altoufaily, Sébastien Baillieul, Benjamin Lemasson, Claire Rome, Anne Briançon-Marjollet

## Abstract

Stroke is a prevalent chronic disease, significantly contributing to mortality and long-term disability. The cerebrovascular recovery process after stroke can be complicated by certain comorbidities, including obstructive sleep apnea syndrome (OSA). However, the mechanisms underlying this deleterious impact of OSA on post-stroke recovery remain elusive. We conducted a preclinical study in rats submitted to stroke and intermittent hypoxia (IH), the main characteristic of OSA, to investigate the effects of IH on stroke lesion and to decipher the pathophysiological mechanisms of this stroke-OSA interaction.

Using a malonate model of ischemic stroke on Sprague Dawley rats exposed to either normoxia or intermittent hypoxia for over 56 days, we monitored brain lesion size, microvascular plasticity and blood brain barrier (BBB) permeability using *in vivo* 4.7T-MRI. Finally, we assessed oxidative stress, inflammation, and angiogenesis-related gene expression by qPCR, and NeuN, GFAP, Collagen IV and ZO-1 expression by immunohistological staining.

Our findings indicate that while IH does not significantly affect lesion volume reduction over time, it exacerbates ischemic injury-induced necrosis and neuronal loss. Additionally, IH amplifies post-stroke inflammation, as evidenced by increased IL-6 and TGF-β expression, and induces oxidative stress by increasing DHE staining and decreasing Superoxide Dismutase (SOD1) level. Vascular assessment revealed IH-induced modifications in vessel radius and angiogenic factors expression (VEGF, Ang2), alongside increased BBB permeability and altered expression of aquaporin 1 and claudin 1, particularly in the acute phase post-stroke.

These results suggest that IH alters cerebrovascular integrity and exacerbates inflammatory response following ischemic stroke, potentially contributing to poorer long-term functional outcomes. Understanding the mechanisms through which IH exacerbates post-stroke injury may guide the development of targeted neuroprotective strategies to improve stroke recovery in patients with OSA.

## Introduction

Stroke is one of the most frequent chronic diseases, affecting both gender and all socio-economic categories of patients. It is estimated that one out of four people will have a stroke in their lifetime and stroke therefore represents the second cause of death and the first cause of disability in adulthood worldwide, with more than six million deaths per 17 million strokes each year [1]. Above 71% of all strokes originate from an infarction of the brain, named ischemic stroke [2]. The recovery process of stroke is usually long and complicated by other morbidities such as obstructive sleep apnea (OSA).

Obstructive Sleep apnea (OSA), characterized by the occurrence of repetition of apnea during sleep, affects nearly 1 billion people worldwide [3]. Severe OSA (apnea-hypopnea index > 30 events per every single hour of sleep) is diagnosed in up to one third of patients with stroke and is associated with an increased risk of recurrent events and overall mortality, rendering OSA screening mandatory after stroke [4, 5]. Therefore, OSA is supposed to represent two different hits related to stroke disease. First OSA increases the risk of stroke occurrence, second OSA -and particularly the Intermittent Hypoxia (IH) caused by OSA-may impair stroke recovery.

Emerging evidence suggests that post-stroke OSA treatment by Continuous Positive Airway Pressure (CPAP) might be beneficial for brain recovery [6, 7]. However, CPAP was not successful in reversing all cardiovascular and cerebrovascular disorders related to OSA and stroke [8], and diagnosing and treating OSA in the acute setting following stroke is often challenging [9–11]. Pressure support is particularly challenging and discontinued in around 50% of stroke patients after one year [9], rendering mandatory the development of alternative therapeutic strategies targeting the detrimental consequences of OSA on the brain. Thus, the development of acute IH-specific neuroprotection may prevent initial stroke worsening and improve long-term functional outcome.

Intermittent hypoxia (IH) triggers key OSA-related cardiovascular and cerebral pathophysiological mechanisms [12, 13]. In rodents, IH exposure before experimental stroke is associated with an increased infarct size and leads to a worsened score of several behavioral tests [14–16]. IH induces brain inflammation, oxidative stress and apoptosis [12, 17–19]. These could alter cerebrovascular function [20] and blood-brain barrier (BBB) integrity [21, 22] and are thought to impair cognitive function [23, 24]. Therefore, IH might contribute to poorer functional and neurocognitive outcomes following stroke [9]. A recent study supports this hypothesis as only 3 days of IH aggravated early brain injury following hemorrhagic stroke [21]. However, the mechanisms underlying this potential deleterious impact of OSA-related IH on cerebrovascular function post-stroke remain to be determined. Indeed, experimental data documenting the effect of IH on the brain after ischemic stroke are scarce, especially over the long-term.

Therefore, our preclinical study in rodents aimed to decipher the effects of IH on stroke lesion and BBB, and the underlying molecular and cellular mechanisms.

## Materials and Methods

### Animals

All animal procedures were run according to the French regulation with the approval of the local ethical committees and the French ministry of research (authorization #24116-2020012918159424 for GIN and #23039-2019112517102122 for HP2). Anesthesia was induced by inhalation of 5% isoflurane (Abbott Scandinavia AB, Solna, Sweden) in 20% O_2_ in air and maintained with 2-2.5% isoflurane through a facial mask (for surgical and imaging procedures). Body temperature was monitored and maintained at 37.0 ± 0.5°C. Weight was measured regularly during the whole experiment (supplementary figure S1).

### Ischemic injury induction

66 male Sprague Dawley rats obtained from Janvier labs weighing 280-300 g (7 weeks old) underwent stereotactic surgery at D0 to induce brain ischemic injury by malonate injection. After local intradermal injection of 0.1 mL of lidocaine 2% and scalp incision, a small hole was drilled (1 mm diameter). Malonate (2 µl, 3M) was injected using a 5 μl-syringe (Hamilton 700 Series, Phymep, Paris, France, 250 μm gauge needle) connected to a micro-pump (injection rate: 0.33 μL/min).The stereotaxic coordinates targeted, were: AP −1.5 mm, ML +3.2 mm, DV −5.3 mm [25] targeting the striatum. After the injection, the needle was left in place for an additional 4 minutes to avoid backflow. The burr hole was sealed, skin resewn and rats were allowed to recover in their home cages. A modified Neurology Severity Score (mNSS) test adopted from [26] was performed 2 hours after the surgery to ensure the succession of the ischemic injury and then at D04, D07, D14 and D28 (supplementary figure S2). One rat died during surgery, thus only 65 rats were included for the rest of the experiment. Due to technical issues such as problems during MRI acquisition or tissue engineering, at some time points the number of rats included for MRI analysis, molecular biology and immunohistology may differ from the total number of rats; but all available samples were always included at each time point. Overview of the number of rats, MRI timepoints and euthanasia timeline in the experimental groups of rats is indicated in Supplementary Table S1.

### Intermittent hypoxia exposure

Starting at D2 after surgery, the rats showing the expected lesion (according to localization, size, NSS) were randomized over lesion size to be exposed to IH or N until D04 (n=10, 5 N and 5 IH), D07 (n=9, 5N and 4 IH), D14 (n=16, 8 N and 8 IH), D28 (n=16, 8 N and 8 IH), and D56 (n=14, 7 N and 7 IH) as described earlier [27]. Briefly, intermittent hypoxia was generated by intermittent injection of low oxygen air in the cages, with 1-min cycles (30 s at 5% FiO_2_, 30 s at 21% FiO_2_ in the cages) repeated for 8 h/day (from 2:00 pm to 10:00 p.m.). Normoxic rats were exposed to the same air flow turbulence and noises as IH rats.

### *In vivo* MRI Experiments

MRI sessions (4.7T, Bruker Avance III, MRI facility of Grenoble IRMaGe) were set at D1, D04, D07, D14, D28 and D56 after malonate injection. Physiological parameters were monitored during MRI acquisition, including body temperature that was maintained at 37°C with help of a warming system. Breath rate was maintained constant (45-60 breath per minute) by modulating inhaled anesthesia. After shimming, MRI sequences were performed as follows: T2 weighted (T2W) images (TR)/echo-time (TE) = 2200/36 ms, voxel size = 117×117×800 μm^3^, 19 slices, field of view (FOV) = 30×30 mm^2^) was performed to determine lesion volume. Diffusion-weighted, spin-echo-EPI (EPI-diff) (TR = 2.2 s, TE = 33 ms, five slices, FOV = 30×30 mm^2^, NA= 8, matrix = 128×128 and voxel size = 234×234×800 μm^3^) was acquired. This sequence was applied 6 times: three times without diffusion weighting and three times with diffusion weighting (b = 800 s.mm^−1^) in three orthogonal directions.

Blood volume fraction (BVF) and Vessel Radius (R) imaging were performed using a steady-state approach. Briefly, multiple gradient-echo sampling of the free induction decay and spin-echo (MGESFIDSE), (TR = 4000 ms, 34 gradient-echoes between 2.3 and 92.3 ms, Spin-Echo = 60 ms; voxel size = 234×234×800 μm^3^, 5 slices, FOV = 30×30 mm^2^) were acquired before and after the manual injection of the UltraSmall SuperParamagnetic Iron Oxide nanoparticles (USPIO®, P904, Guerbet, France; 133 μmo lFe/kg). A three-minute delay after the first injection was applied before starting the second MGEFIDSE acquisition.

BBB leakage was assessed using the dynamic contrast enhanced (DCE) sequence involved 25 images of T1-weighted (T1W) spin-echo images (TR/TE = 800/4 ms, voxel size = 234×234×800 μm^3^, 5 slices, FOV = 30×30 mm^2^). The acquisition time of one repetition was 15.2 s. After the acquisition of four baseline images, a bolus of Gd-DOTA (Dotarem®, Guerbet, France; 200 μmol/kg) was administered through the tail vein. Perfusion-related parameters as well as TTP and percentage of signal enhancement were measured on the DCE signal curve during the first 3 min post-injection of the bolus.

### MRI Data Processing

All processing was computed, pixel by pixel, using a home-made program developed using MATLAB (MathWorks, Natick, MA, USA).

ADC maps were automatically computed on the Bruker scanner as the means of the ADCs observed in each of three orthogonal directions. BVF, R and SO_2_ were computed using the MR fingerprinting approach proposed by Delphin et al. [28]. Note that this dictionary was simulated for experiments using USPIO with a 200 μmol Fe/kg dose. BVF, R and SO_2_ reflect only the functional vessels. BBB permeability was calculated on the DCE images as the signal enhancement (SE, %) induced by Gd-DOTA extravasation.

For lesion volume, the region of interest (ROI) corresponding to the whole ischemic lesion was manually delineated on T2W images. The same volume was also delineated on the contralateral hemisphere as a control. Viable ROI was defined as a sub-region of the lesional ROI using these constraints: ADC = [0–2500], R > 0; BVF > 0 and SO_2_ = [0 – 100 %]. Viable ROI was defined as a sub-region of the lesional ROI using these constraints: ADC = [0-2500 µm^2^.s^−1^], R >0 µm; BVF >0 % and StO_2_ = [0-100 %]. The remaining non-viable part of the lesion is referred to as necrotic ROI. After extracting the brain using the PCNN3D algorithm [29], T_2_W anatomical images and lesion regions of interest were co-localised using the FLIRT function of FSL (the FMRIB Software Library [30]) for all animals imaged at D1 (n=33), and finally all co-localized lesions were added.

### RT-qPCR

At the end of the exposure, animals were anesthetized with isoflurane then euthanized by decapitation, and brains were collected and frozen at −80°C. To measure the expression level of genes of interest at D04, D07 and D28, reverse transcription-quantitative polymerase chain reaction (RT-qPCR) experiments were done. Each brain section was divided into two hemisections where the contralateral hemisphere served as a control for each lesioned hemisphere (ipsilateral). Total RNA was extracted from frozen hemisections with TRIzol® reagent (Invitrogen, Life Technologies Ltd., UK) and total RNA quantification was performed on a spectrophotometer (Nanodrop ND2000, Thermo Fisher Scientific, USA). Reverse transcription was performed under the conditions recommended by the manufacturer (iScript, Biorad, USA). Quantitative PCR was performed on thermocycler® (C1000, BioRad, USA) using the SYBRGreen method (SsoAdvanced Sybr green, BioRad, USA) and primer pairs (Sigma Aldrich Merck, Germany) referenced in Supplementary Table S3. The specificity of each PCR product was assessed using the dissociation reaction plot. Relative gene expression was calculated with the 2^−ΔΔCT^ method, using the level of Cyclin A (*CYCA*) as a normalization gene. Samples were duplicated for each gene analysis.

### Immuno-histological staining

At D04, D07 and D28, immunohistology was performed on 10 µm frozen brain sections. Sections were dropped on Superfrost™ Excell™ adhesion microscope slides, fixed with PFA 4%, permeabilized with PBS tween 0.1% BSA 3% Triton 0.5%, and blocked with 2% BSA, then incubated overnight at 4°C with primary antibodies (supp. table 1) in a dark moist chamber, washed with PBS, and incubated 2 h at room temperature with secondary antibodies (see Supplementary Table S2) and DAPI (1 μg/mL). We characterized vessels and BBB (Collagen IV, ZO-1), number of neurons (NeuN) and astrocytic activation (GFAP). Images were obtained using an epifluorescence microscope (AxioScan, Zeiss, Germany) on PIC-GIN platform (https://neurosciences.univ-grenoble-alpes.fr/fr/picgin).

### Immuno-histological analysis

Data were analyzed using QuPath. For GFAP, ZO1 and ColIV, a random trees pixel classifier was used for analysis, this classifier is able to detect the surface staining per pixel in the total surface area of our region of interest (the viable part of the lesion). For NeuN, an object classifier was employed to detect co-stained DAPI cells and NeuN-positive cells. During analysis, 2-3 images per rat were analyzed and each rat served as its own control by comparing lesioned hemisphere (ipsilateral) to contralateral hemisphere.

### Dihydroethidium (DHE) staining

DHE staining was performed at D04 and D07. The staining was adapted from [31] with the following modifications: frozen brain samples were cryosectioned at 10 μm thick and allowed to air dry for 15 min at room temperature onto Superfrost™ Excell™ adhesion microscope slides (Dutscher, France). DHE 10 μM (Sigma-Aldrich Merck, Germany) was added onto brain sections for 5 minutes at room temperature (PBS was used as control in contralateral sections) in a dark moist chamber. After washing, cover slips were added and the fluorescent signal was recorded using microscopy (CKX53SF, Olympus, Evident, Germany) and analyzed with ImageJ software (https://imagej.net/ij/). DHE-positive cells in the perilesional area were expressed as the percentage of brain area and normalized to the contralateral hemisphere.

### Hematocrit measurements

Blood was collected under gas anesthesia with isoflurane (2%) from the caudal vein, in vacutainer lithium-heparin tubes, after 2 hours of fasting, and stored on ice before being rapidly centrifuged at 1600g for 15 minutes at 4°C. After blood collection, hematocrit measurements were done in the plasma at D07, D28 and D56 (supplementary figure S3).

### Statistical analysis

For MRI, weight, lesion volume, mNSS, and hematocrit measurements, two-way Anova was used on GraphPad Prism software. For gene expression and immunohistological analysis, three-way Anova was used on Statistica software. Results are represented with a median with interquartile range. Any significant result was considered when p value < 0.05. When a significant interaction was found between IH, malonate and time in the three-way Anova, Tukey HSD post-hoc tests were performed and p-values for differences between time-points in IH were reported in Supplementary Table S4.

## Results

### Intermittent hypoxia does not affect lesion volume after ischemic injury

Lesion volume (mm³) was measured on T2W images obtained from non-invasive MRI scans at D1, D07, D14, D28, and D56 following the induction of focal ischemia (Figure 1A, 1B and [32]). The median lesion volume after malonate injection significantly decreased from 47 mm³ for N group and 43 mm³ for IH group at D1, 33 mm³ for N group and 24 mm³ for IH group at D04, to 15 mm³ for N group and 18 mm³ for IH group at D07 (p = 0.0002)(Figure 1A). The lesion volumes had further decreased to 12 mm³ in the N group and 12 mm³ in the IH group (p < 0.0001). At D28, the lesion volumes were 11 mm³ in the N group and 12 mm³ in the IH group (p < 0.0001), and at D56, they were 10 mm³ in the N group and 14 mm³ in the IH group (p < 0.0001). These findings indicate a significant recovery phase within the first 14 days post-lesion. However, no significant changes in lesion volume were observed in the IH group compared to N (Figure 1A).

**Figure 1.**
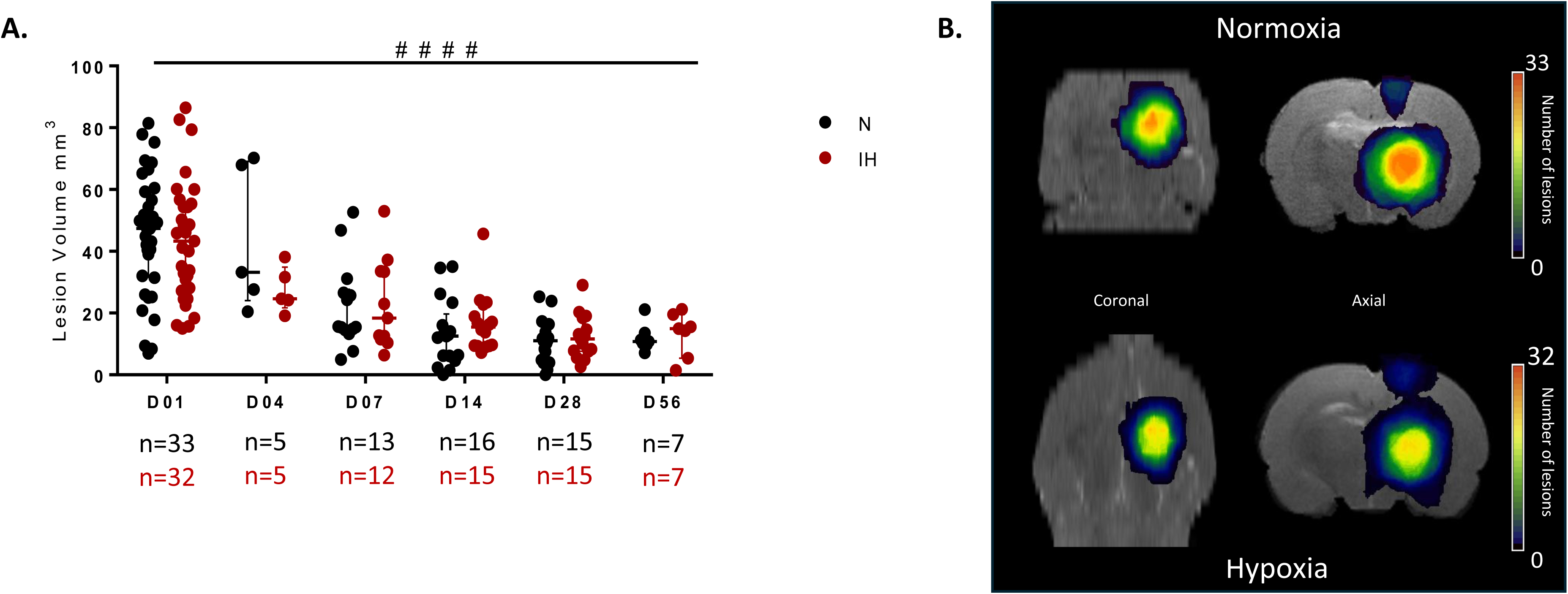
IH does not affect brain lesion volume after 4, 7, 14, 28 and 56 days post ischemic injury induced by malonate injection. A) Median lesion volume at different time points after IH/N ischemic injury showing that lesion size significantly decreased from its original size starting from D07 until D56 with no effect of IH. # indicates the effect of time. The number of animals at each time point is indicated under the figure: at D01 n=33 in N and 32 in IH, at D04 n=5 in each group, at D07 n=14 N and 11 IH, at D14 n=17 N and 15 IH, at D28 n=17 N and 16 IH, at D56 n=7 in each group. B) Overlap map with color-coded injured voxels from T_2_W anatomical images and lesion regions of interest at D1 for the normoxic (n=33) and hypoxic groups (n=32), providing an overview of all the lesioned brain areas after injection of malonate. ####: p<0.0001 vs D01 (two-way ANOVA).

### Intermittent hypoxia exacerbates necrosis and neuronal loss induced by ischemic injury

To assess the level of necrosis within the lesion, we calculated from T2W MRI images the apparent diffusion coefficient (ADC, µm²/sec) (Figure 2A), and we also calculated the necrotic volume % from the total lesion volume (Figure 2B). In addition, neuronal loss (NeuN) was characterized by immunohistological analysis on brain sections after 4, 7 and 28 days of ischemic injury and IH/N exposure (Figure 2C, 2D). Necrosis increased with time in the lesioned hemisphere compared to the contralateral hemisphere (Figure 2A and 2B), and the number of viable neurons decreased (NeuN+ cells)(Figure 2C). Moreover, compared to normoxia, IH further increased tissue necrosis induced by ischemic injury on D28 (Figure 2A). IH also further decreased the number of viable neurons (Figure 2C).

**Figure 2.**
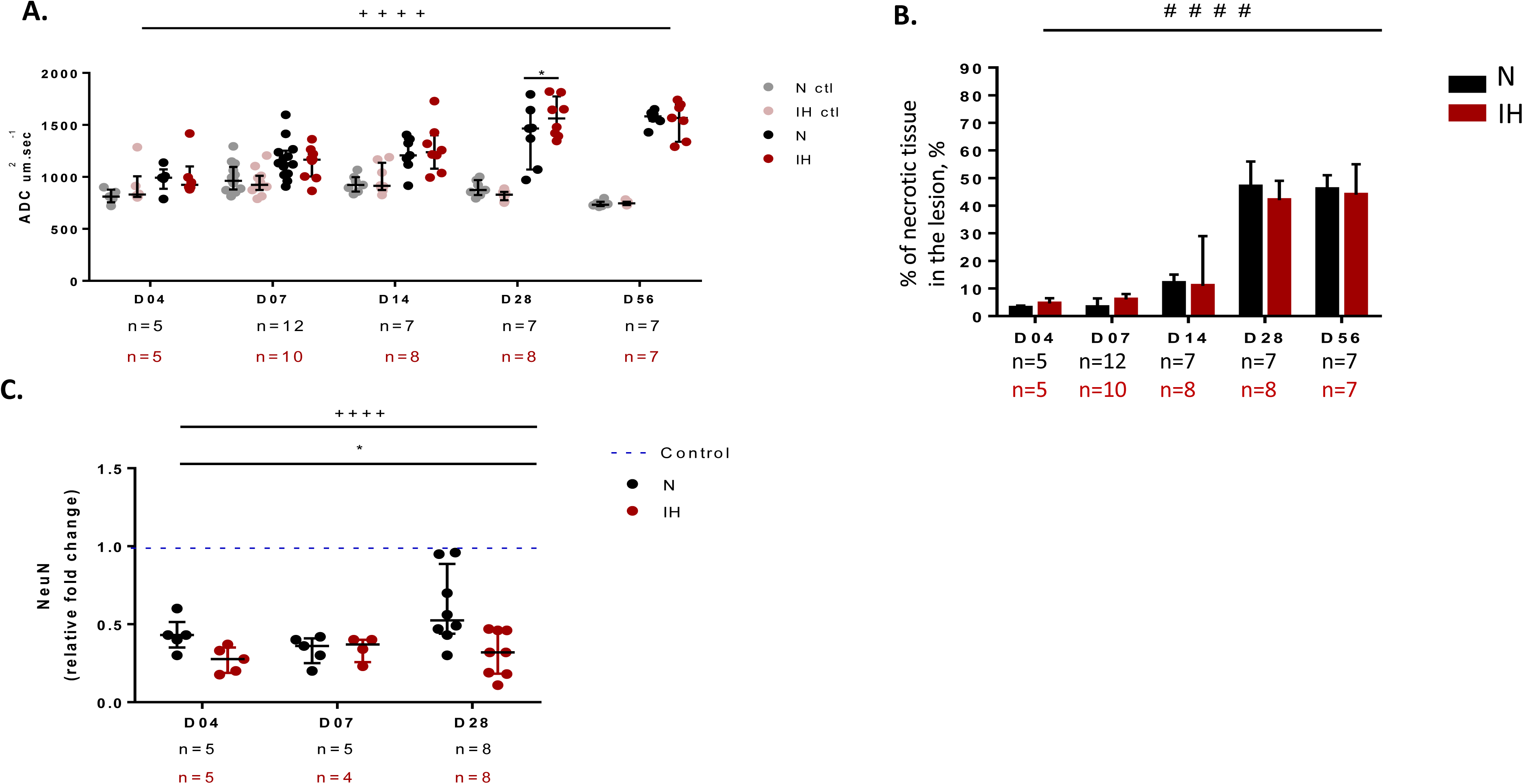
IH exacerbates tissue necrosis after ischemic injury. A) Evolution of ADC calculated from MRI T2W images. N control and IH control corresponds to the contralateral hemisphere, N and IH to the ipsilateral (lesioned) hemisphere. B) Percentage of necrotic tissue with respect to the total lesion volume. C) Relative expression of NeuN-positive cells in the viable part of the lesion. The values were normalized to the contralateral hemisphere referred to as control. Data is represented by median with interquartile range. */+: 0.03>p>0.01, ****/++++: p<0.0001 three-way ANOVA. * indicates the difference between IH and N, + indicates the effect of malonate (ipsilateral vs contralateral hemisphere). The number of animals at each time point is indicated under the figure: D04 n=5 in each group, D07 n=10-12, D14 n=7-8, D28 n=7-8, D56 n=7 in each group. NeuN: neuronal nuclear antigen, ADC: apparent diffusion coefficient, ROI: region of interest, ctl : contralateral.

### Intermittent hypoxia exacerbates inflammation induced by ischemic injury

The expression levels of genes associated with inflammation (IL-6, TGF-β, TNF-α, NF-κB) were tracked over time (D04, D07, and D28) through qPCR at the site of the ischemic lesion after N/IH exposure (Figure 3A). In addition, to characterize astrocytic activation (GFAP), immunohistological analysis was performed on brain sections after 4, 7 and 28 days of ischemic injury after N/IH exposure (Figure 3B, 3C). The results revealed increased expression of IL-6, TGF-β and GFAP in the lesioned hemisphere compared to the contralateral hemisphere. Furthermore, in the IH group, there was a further increase in the inflammatory markers IL-6 and TGF-β compared to normoxia, with no significant effect on TNF-α or NF-κB (Figure 3A). IH exposure also led to significant increase in astrocytic activation (GFAP+ cells) compared to N condition (Figure 3B).

**Figure 3.**
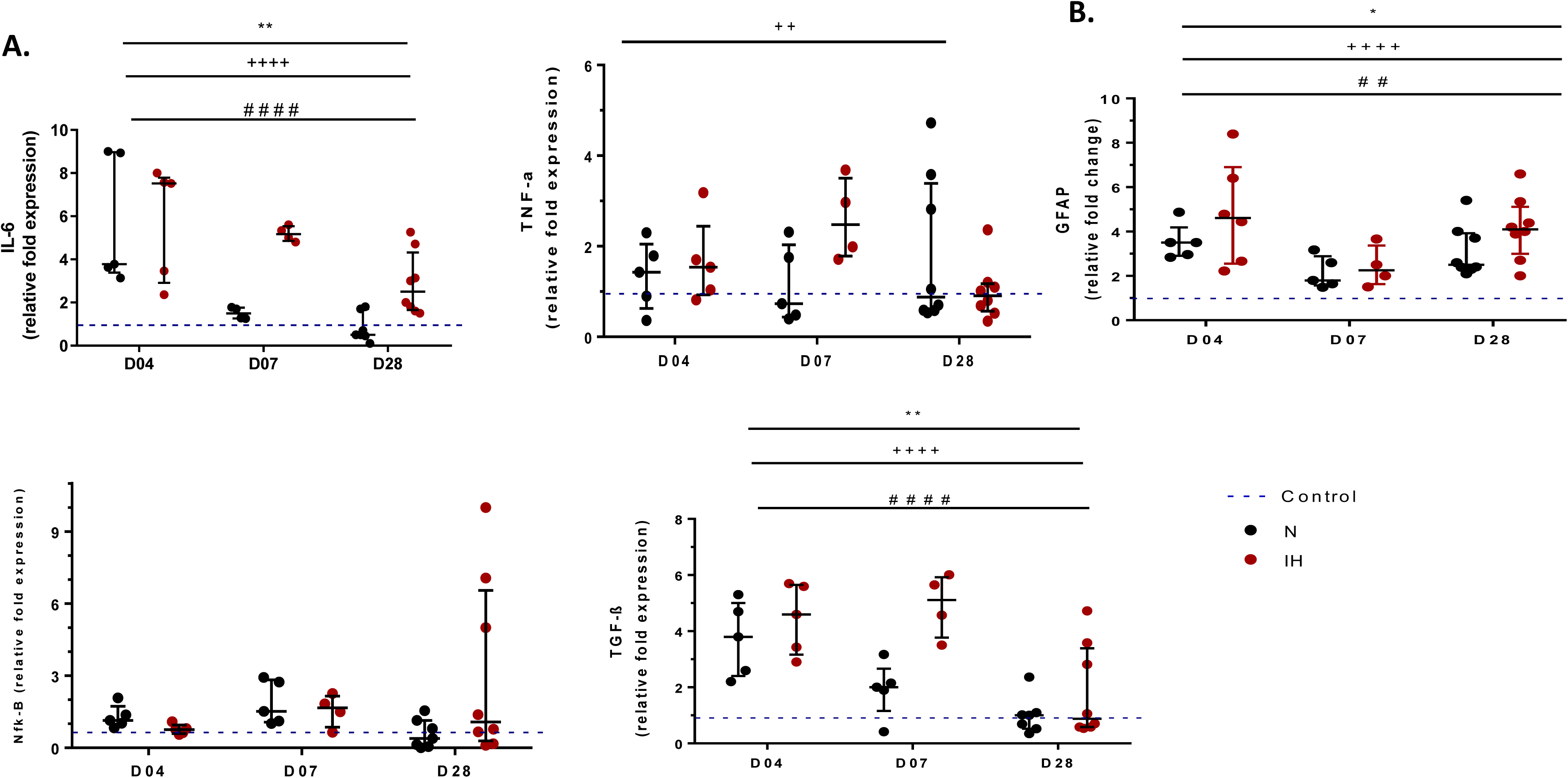
IH exacerbates inflammation after ischemic injury. A) Relative brain gene expression of IL-6, TGF-β, TNF-α, NF-κB at D04 (n=4-5 per group), D07 (n=4-5 per group), and D28 (n=7-8 per group). B) Relative brain expression of GFAP surface staining per pixel in the total surface area of our region of interest (the viable part of the lesion) at D04 (n=5-6 per group), D07(n=5 per group) and D28 (n=8 per group). The values were normalized to the contralateral hemisphere referred to as control. Data is represented by median with interquartile range. */#/+: 0.03>p>0.01, ***/###/+++: 0.0005>p>0.0001, ****/####/++++: p<0.0001, three-way ANOVA. * indicates the difference between IH and N, + indicates the effect of malonate (ipsilateral vs contralateral hemisphere), # indicates the difference between days. GFAP: glial fibrillary acidic protein, IL-6: interleukin 6, NF-kB: nuclear factor kappa light chain enhancer of activated B cells, TNF-alpha: tumor necrosis factor alpha, TGF-B: transforming growth factor beta.

### Intermittent hypoxia induces oxidative stress in the lesioned hemisphere

The expression levels of genes associated with oxidative stress (SOD1, catalase, Nrf1 and HIF-1α) were tracked over time (D04, D07, and D28) through reverse transcription-quantitative PCR (qPCR) on the total hemisphere of the ischemic lesion (Figure 4A). In addition, dihydroethidium (DHE) staining was performed at D04 and D07 to monitor superoxide production (Figure 4B, 4C). Malonate injection induced an increase of Nrf1 gene expression, without changes in DHE staining or SOD expression. By contrast, compared to normoxia, IH increased DHE in the perilesional area at D04 (Figure 4B) and decreased antioxidant enzyme SOD1 at all time points (Figure 4A) without affecting other parameters.

**Figure 4.**
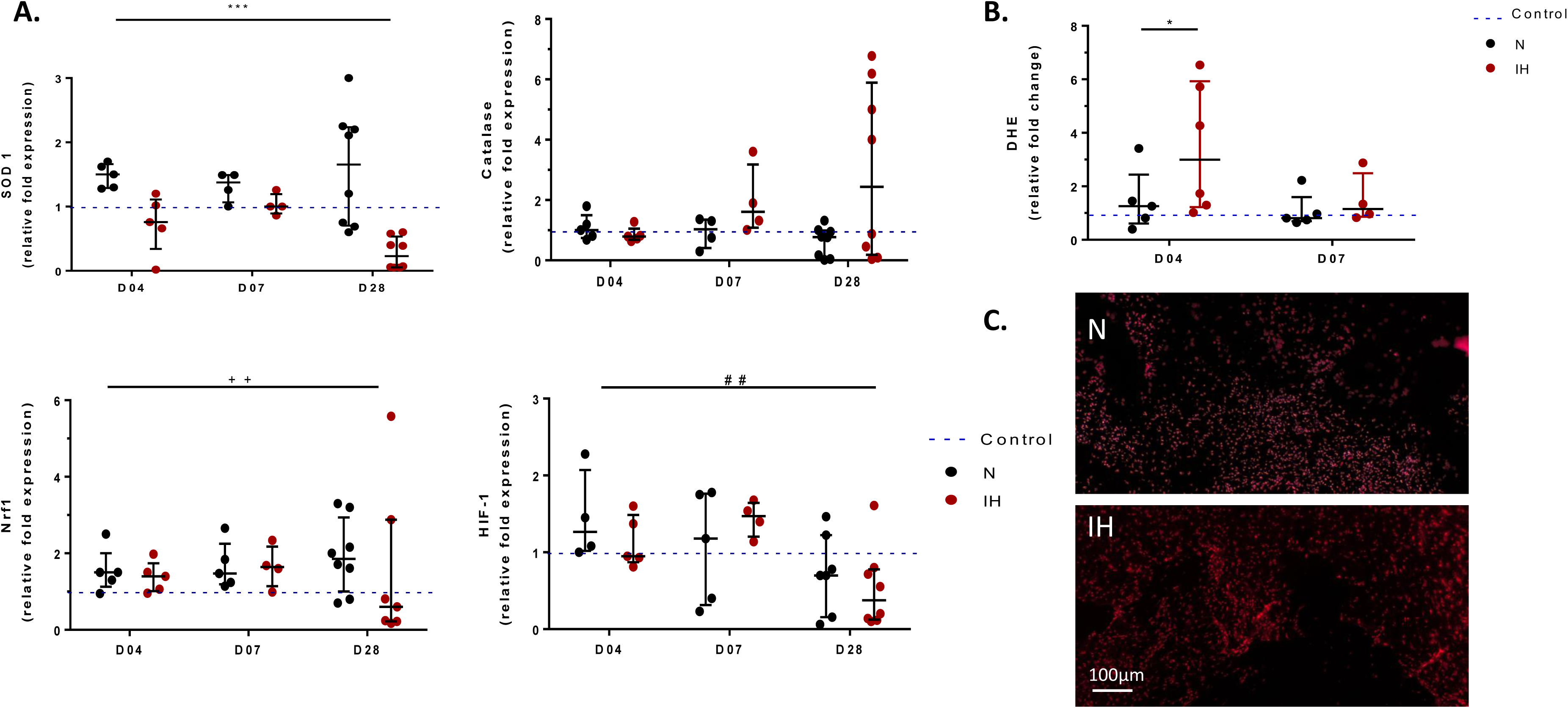
IH exacerbates oxidative stress after ischemic injury. A) Relative brain gene expression of SOD1, catalase, NRF1, and HIF-1 at D04 (n=4-5 per group), D07(n=4-5 per group), and D28 (n=7-8 per group). B) Relative expression of DHE-positive cells in the perilesional area expressed as the percentage of stained brain area at D04 (n=5-6 per group) and D07 (n=4-5 per group). The values were normalized to the contralateral hemisphere referred to as control. C) Representative image of DHE staining at D04 in the perilesional area of N and IH rats. Data is represented by median with interquartile range. */#/+: 0.03>p>0.01, ***/###/+++: 0.0005>p>0.0001, three-way ANOVA. * indicates the difference between IH and N, + indicates the effect of malonate (ipsilateral vs contralateral hemisphere), # indicates the difference between days. DHE: dihydroethidium, Nrf1: nuclear respiratory factor 1, SOD1: superoxide dismutase 1.

### Intermittent hypoxia alters brain vasculature after ischemic injury

To assess the effects of IH after ischemic injury on blood volume (BVF), vessel radius (R) and tissue oxygenation (StO_2_), a multi-gradient sequence was performed two minutes before and after the intravenous injection of intravascular (IV) USPIO particles (P904®). We measured blood volume flow (%), vessel radius (µm), and oxygen saturation (%) at 4, 7, 14, 28, and 56 days following malonate injection and IH/N exposure (Figure 5A). In addition, the expression levels of genes associated with growth factors and angiogenesis (VEGF, VEGFR1, VEGFR2, Ang1, and Ang2) were tracked over time (D04, D07, and D28) on the total hemisphere of the ischemic lesion (Figure 5B). Finally, extracellular matrix component ColIV was characterized by immunohistological analysis done on brain sections after 4, 7 and 28 days of ischemic injury and IH/N exposure (Figure 5C, 5D). MRI analysis showed that malonate injection profoundly impacted vessel radius (R), blood volume (BVF) and O_2_ saturation in the brain, as well as gene expression for VEGF, VEGRs and Ang1 and 2. Moreover, compared to normoxia, IH increased vessel radius at D28 (Figure 5A). Brain characterization showed that compared to normoxia, IH increased the gene expression of VEGF, VEGFR1 and decreased the expression of Ang2 at D07. However, IH then decreased VEGF, VEGFR1 and VEGFR2 and increased Ang2 at D28. Finally, we observed significant differences between consecutive days (IH D04 vs. D07 vs D28), indicating a time-dependent effect of intermittent hypoxia and malonate on VEGF, VEGFR1, VEGFR2 and Ang2 (See supplementary table 3).

**Figure 5.**
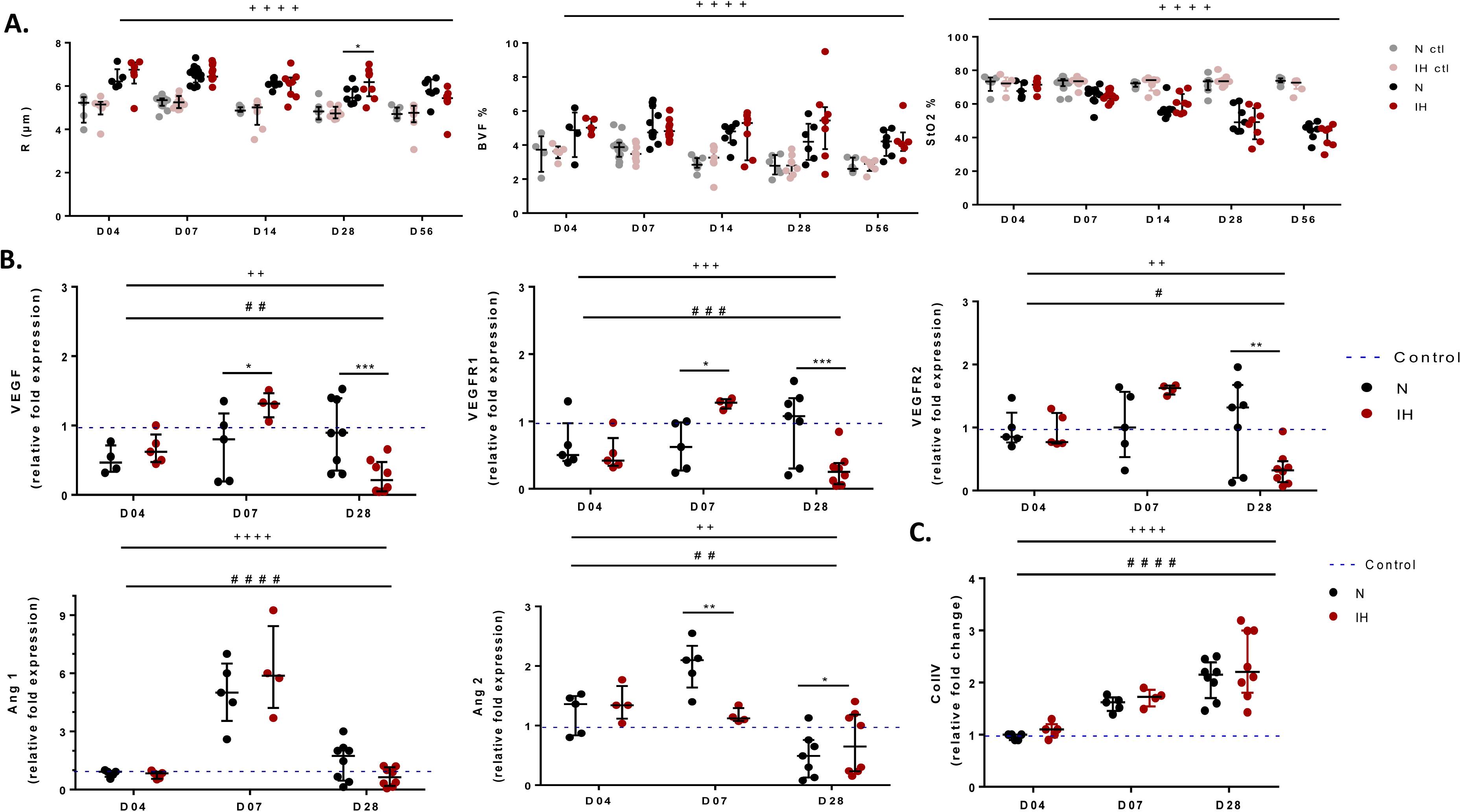
IH alters brain vasculature after ischemic injury. A) Vessel radius (R), Blood volume (BVF), and oxygen saturation (StO_2_) with their corresponding maps generated by MR fingerprinting approach, at D04 (n=4-5 per group), D07 (n=10-12 per group), D14 (n=7-8 per group), D28 (n=7-8 per group), and D56 (n=7 per group). B) Relative brain gene expression of VEGF, VEGFR1, VEGFR2, Ang1, and Ang2 at D04 (n=4-5 per group), D07 (n=4-5 per group), and D28 (n=7-8 per group). C) Relative brain expression of Collagen IV surface staining per pixel in the total surface area of our region of interest (the viable part of the lesion area) at D04 (n=5-6 per group) and D07 (n=4-5 per group) and D28 (n=7-8) per group. The values were normalized to the contralateral hemisphere referred to as control. Data is represented by median with interquartile range. */#/+: 0.03>p>0.01, ***/###/+++: 0.0005>p>0.0001, ****/####/++++: p<0.0001, three-way ANOVA. * indicates the difference between IH and N, + indicates the effect of malonate (ipsilateral vs contralateral hemisphere), and # indicates the difference between days. Ang1/2: angiopoietin 1 and 2, VEGF: vascular endothelial growth factor, VEGFR1/R2: vascular endothelial growth factor receptor 1 and 2.

### Intermittent hypoxia exacerbates blood-brain barrier permeability after ischemic injury

The effect of IH on BBB permeability after ischemic injury was calculated as the signal enhancement (%) induced by gadolinium extravasation (Figure 6A). In addition, tight junction ZO-1 was characterized by immunohistological analysis done on brain sections after 4, 7 and 28 days of ischemic injury and IH/N exposure (Figure 6B), and the expression levels of genes associated with BBB integrity (occludin, claudin 1, claudin 5, Aquaporin 1, MRP1, and P-gp) were tracked over time (D04 and D07) on the total hemisphere of the ischemic lesion (Figure 6C). Compared to normoxia, the BBB was more permeable after IH on D04, without a significant effect on other time points (Figure 6A). Moreover, while malonate injection increased claudin 1 and aquaporin 1 expression at D07 in normoxic rats, these expressions were decreased in IH compared to N, without affecting other tight junction proteins (claudin 5 and occludin) or transporters (P-gp and MRP1). Finally, IH decreased the level of ZO-1 at D28 compared to N.

**Figure 6.**
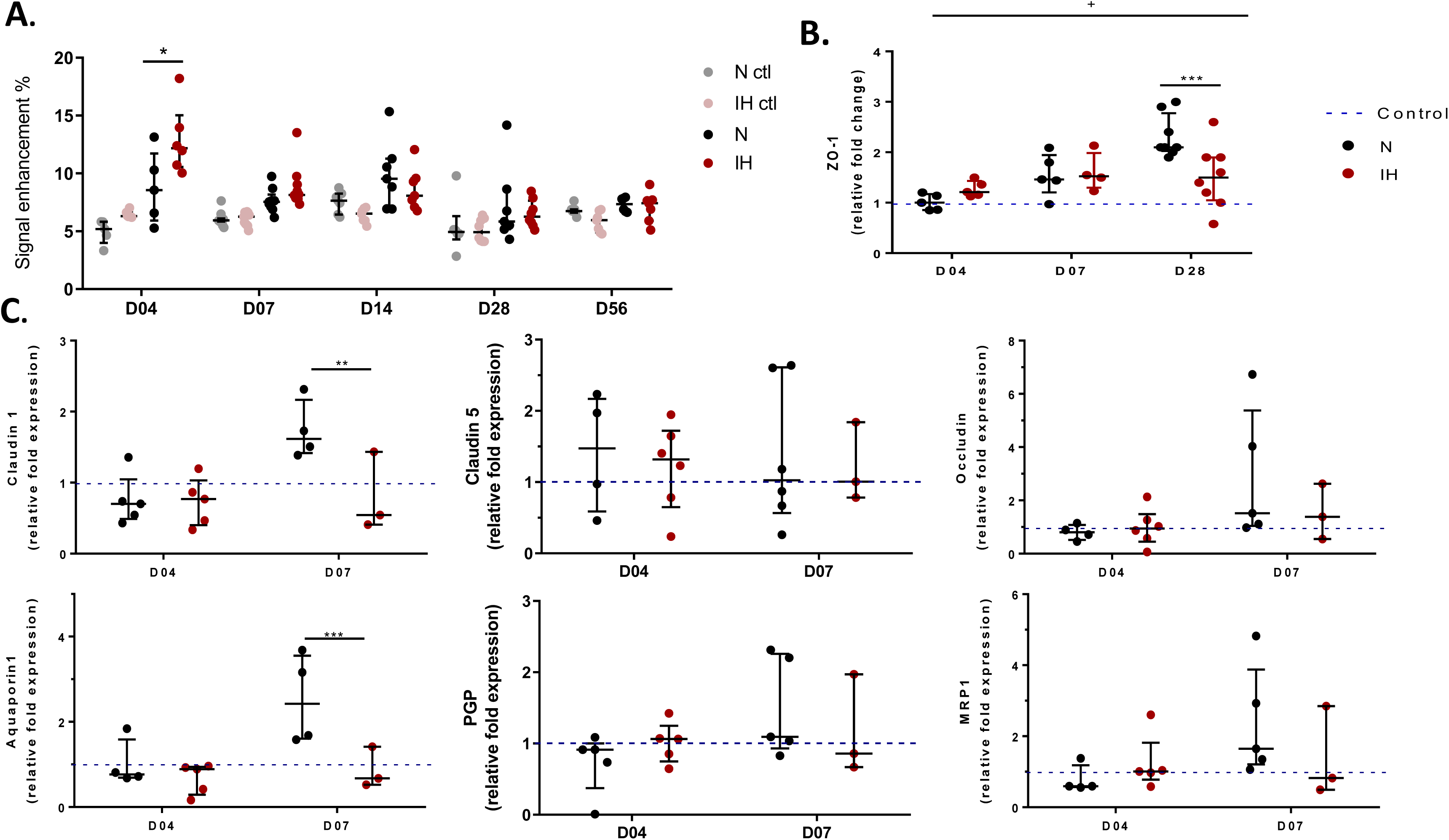
IH exacerbated BBB permeability after ischemic injury. A) BBB permeability measured by signal enhancement on MRI at D04 (n=4-5 per group), D07 (n=7-8 per group), D14 (n=7-8 per group), D28 (n=7-8 per group), and D56 (n=7 per group). B) Relative brain expression of ZO-1 surface staining per pixel in the total surface area of our region of interest (the viable part of the lesion area) at D04 (n=5-6 per group) and D07 (n=4-5 per group) and D28 (n=7-8) per group. The values were normalized to the contralateral hemisphere referred to as control. C) Relative brain gene expression of claudin 1, claudin 5, occludin1, aquaporin 1, MRP1 and P-gp at D04 (n=3-5 per group), and D07 (n=3-5 per group). Data is represented by median with interquartile range. */#/+: 0.03>p>0.01, ***/###/+++: 0.0005>p>0.0001, three-way ANOVA. * indicates the difference between IH and N, + indicates the effect of malonate (ipsilateral vs contralateral hemisphere), and # indicates the difference between days. MRP1: Multidrug resistance-associated protein 1, P-GP: P-glycoprotein, ZO-1: zonula occludens 1.

## Discussion

This study aimed to investigate the impact of intermittent hypoxia on cerebrovascular recovery following ischemic injury. Our findings suggest that while IH does not significantly affect lesion volume reduction as measured by MRI over time, it exacerbates secondary injury mechanisms, including inflammation, oxidative stress, alterations in vascular remodeling and blood-brain barrier disruption. We employ the malonate-induced focal ischemia model, a well-established approach for studying ischemic stroke, as previously characterized in experimental stroke research and fully described in a previous study [32]. The malonate model offers a controlled and reproducible method for inducing ischemic-like neuronal injury. This model provides a reliable platform for investigating key pathophysiological mechanisms underlying stroke, including oxidative stress, inflammation, and neurovascular dysfunction. Building upon previous findings, we utilize this model to explore the impact of intermittent hypoxia (IH), a hallmark of obstructive sleep apnea (OSA), on post-stroke cerebrovascular injury.

MRI analysis revealed a significant decrease in lesion volume over time, particularly within the first 14 days following ischemic injury, indicating an initial recovery phase. However, exposure to IH did not significantly alter the lesion size as measured by MRI at any time point, suggesting that IH does not directly influence infarct resolution. This contrasts with findings from other studies suggesting that IH exposure before stroke could increase infarct size [14–16]. Even though these studies were performed with occlusion models while ours uses the malonate stroke model, these findings suggest different impacts of IH when it is applied before or after the experimental stroke.

As expected in a stroke model, malonate injection induces necrosis accompanied by inflammation and moderate oxidative stress in the lesioned hemisphere (Figure 2 to 4 and [32]). We found that IH exacerbated these phenomenons with further increased necrosis, decreased number of viable neurons (NeuN), increased inflammation marker IL-6, increased astrocytic reaction (GFAP), and increased oxidative stress (elevated DHE staining accompanied with altered SOD expression) throughout the longitudinal follow-up. These findings are in line with previous data showing that IH induced inflammation, oxidative stress and apoptosis in the brain, independently of stroke [12]. In particular, an increase of IL-6 [17, 33] and GFAP [34, 35] after IH exposure were documented along with classical inflammatory markers such as TNF-ɑ or iNOS [12]. Similarly, several studies report an elevation of DHE staining[36, 37] and a diminution of the antioxidant enzyme SOD expression [12] in the brain after IH exposure, along with other markers or enzymes such as MDA and NADPH oxidase [12]. Interestingly, malonate injection also increased the expression of the anti-inflammatory cytokine TGF-β, suggesting the initiation of an anti-inflammatory response in early stages after stroke (Figure 3 and [32]). While one study reported a decrease of TGF-β in IH-exposed mice brains [38], interestingly in our study IH after stroke further increased TGF-β expression, suggesting an enhanced effect on both pro- and anti-inflammatory responses after IH. Our findings thus confirm the known impact of IH on brain inflammation and oxidative stress and expand it to the context of post-stroke recovery. Since inflammation and oxidative stress are associated with brain lesion after ischemic stroke in both occlusion and malonate models [32, 39, 40] and their exacerbation by IH after stroke may participate in delaying proper recovery.

Inflammation after ischemic stroke is associated with a transient alteration of BBB permeability characterized by overexpression of aquaporin 1 and claudin 1 at D07, and later expression of ZO-1 at D28 that leads to BBB stabilization through tight junctions (Figure 6 and [32]). Herein, we observe that IH increases BBB permeability in the acute phase after stroke, as evidenced by MRI at D04. IH also abolishes malonate-induced changes in claudin 1, Aquaporin 1 and ZO-1. The decrease in ZO-1 and AQP1 under IH confirms previous findings in a cellular BBB model [41] and *in vivo* [42], where low AQP1 was associated with brain edema. Similarly, Roche et al. [22] observed an increase in Claudin-1 associated with increased BBB permeability after acute (1 day) exposure to IH, which could participate in BBB disruption [43]. In our experiments, the absence of claudin 1 increase under IH at D07 suggests early disruption of BBB, followed by a BBB recovery process.

After malonate injection, we also observe vascular remodeling evidenced by MRI (vessel radius, BVF, StO_2_) as well as by increased expression of Angiopoietin 1 and 2 and collagen IV (Figure 5 and [32]) corresponding to angiogenesis and microvessel structural and functional remodeling after stroke. When exposed to IH, we observe further augmentation of vessel radius at D28, associated with a complex dynamic regulation of vascular growth factors family members VEGF, Ang1 and Ang2. Indeed, at D07 after stroke VEGF signaling pathway genes (VEGF, VEGFR1, VEGFR2) were upregulated compared to normoxia. In parallel, at D07 Ang1 is elevated while IH abolishes the malonate-induced increase of Ang2. By contrast, at D28 IH increases Ang2, while VEGF and VEGFRs expressions are blunted. VEGF, a known pro-angiogenic factor, seems to be elevated in rodent brains after 14 days of IH [44]. Although VEGF expression in the brain was sparsely studied, this increase is consistent with abundant literature documenting similar upregulation of VEGF in OSA/IH models [45–47]. Angiopoietins are members of the vascular growth factor family, Ang1 being considered as a pro-angiogenic and vessel stabilization factor, while Ang2 has a complex role associated with vessel destabilization, angiogenesis and pathological remodeling [48, 49]. Interestingly, Ang2 implication was suggested in IH-induced angiogenesis in the rodent hippocampus [44]. Altogether, our results suggest that IH-induced elevation of VEGF and VEGFR associated with decreased Ang2 at D07 may favor early angiogenesis after stroke. However, at D28 blunted VEGF, increased Ang2 and decreased ZO-1 compared to normoxia suggest that this angiogenic response induced by IH could be only temporary in subacute stages of stroke recovery (D07), and followed by pathological microvascular remodeling at later stages.

In this study, we used a model of focal ischemic injury induced by malonate injection in the striatum. This model was developed recently [25] and further characterized in MRI and with molecular analysis by our team [32]. This model is reproducible and associated with low mortality. It induces a minor, lacunar stroke, and is relevant to mimic stroke related to occlusion of the deep perforating arteries. To further enhance our understanding of the IH impact on the brain after stroke, future studies could consider using the transient Middle Cerebral Artery Occlusion (MCAO) model. This model is less reproducible and associated with higher mortality, but mimics larger strokes as induced by an occlusion of the middle cerebral artery, thus providing additional information related to a different type of stroke.

## Conclusion

This study demonstrates that while IH does not affect lesion volume post-stroke as measured by MRI, it exerts deleterious effects after stroke by exacerbating inflammatory and oxidative stress responses, neuron death and early BBB disruption. In the subacute stage after stroke, IH could favor angiogenesis through VEGF and Ang signalling pathways. However, this temporary response appears to be blunted in later stages after stroke, where inflammation and dysfunctional remodeling principally characterize the brain response to IH. Future studies should explore therapeutic strategies, for example targeting inflammation or oxidative stress, aimed at mitigating the adverse effects of IH to improve post-stroke outcomes.

## Abbreviations and acronyms

ADC: apparent diffusion coefficient
Ang1: angiopoietin 1
Ang2: angiopoietin 2
BBB: blood-brain barrier
BVF: blood volume fraction
ColI-IV: collagen type IV
DAPI: 4′,6-diamidino-2-phenylindole
DHE: dihydroethidium
FOV: field of view
FSL: FMRIB Software Library
GFAP: glial fibrillary acidic protein
HIF-1: hypoxia inducible factor 1
IH: Intermittent hypoxia
IL-6: interleukin 6
IV: intravenous injection
MRI: magnetic resonance imaging
MRP1: multidrug resistance-associated protein 1
NeuN: neuronal nuclear antigen
Nf-kB: nuclear factor kappa light chain enhancer of activated B cells
Nrf1: nuclear respiratory factor 1
OSA: Obstructive Sleep Apnea
PBS: phosphate-buffered saline
P-GP: p-glycoprotein
R: vessels radius
ROI: region of interest
RT-qPCR: reverse transcription-quantitative polymerase chain reaction
SE: signal enhancement
SOD1: superoxide dismutase 1
StO_2_: oxygen saturation
T_1_W: T_1_ weighted
T_2_W: T_2_ weighted
TGF-B: transforming growth factor beta
tMCAO: transient middle cerebral artery occlusion
TNF-alpha: tumor necrosis factor alpha
VEGF: vascular endothelial growth factor
VEGFR1: vascular endothelial growth factor receptor 1
VEGFR2: vascular endothelial growth factor receptor 2
ZO-1: Zona occludens 1

## Acknowledgments

The authors thank the animal facility working teams for their assistance in animal care and housing at the GIN, HP2 and PHTA animal facilities. This work was supported by the Photonic Imaging Center of Grenoble Institute Neuroscience (Univ Grenoble Alpes–Inserm U1216) which is part of the ISdV core facility and certified by the IBiSA label. This work was performed on the IRMaGe platform member of France Life Imaging network (grant ANR-11-INBS-0006) and on the HypE platform member of IBiSA network.

## Supplementary Data

**Supplementary figure 1.**
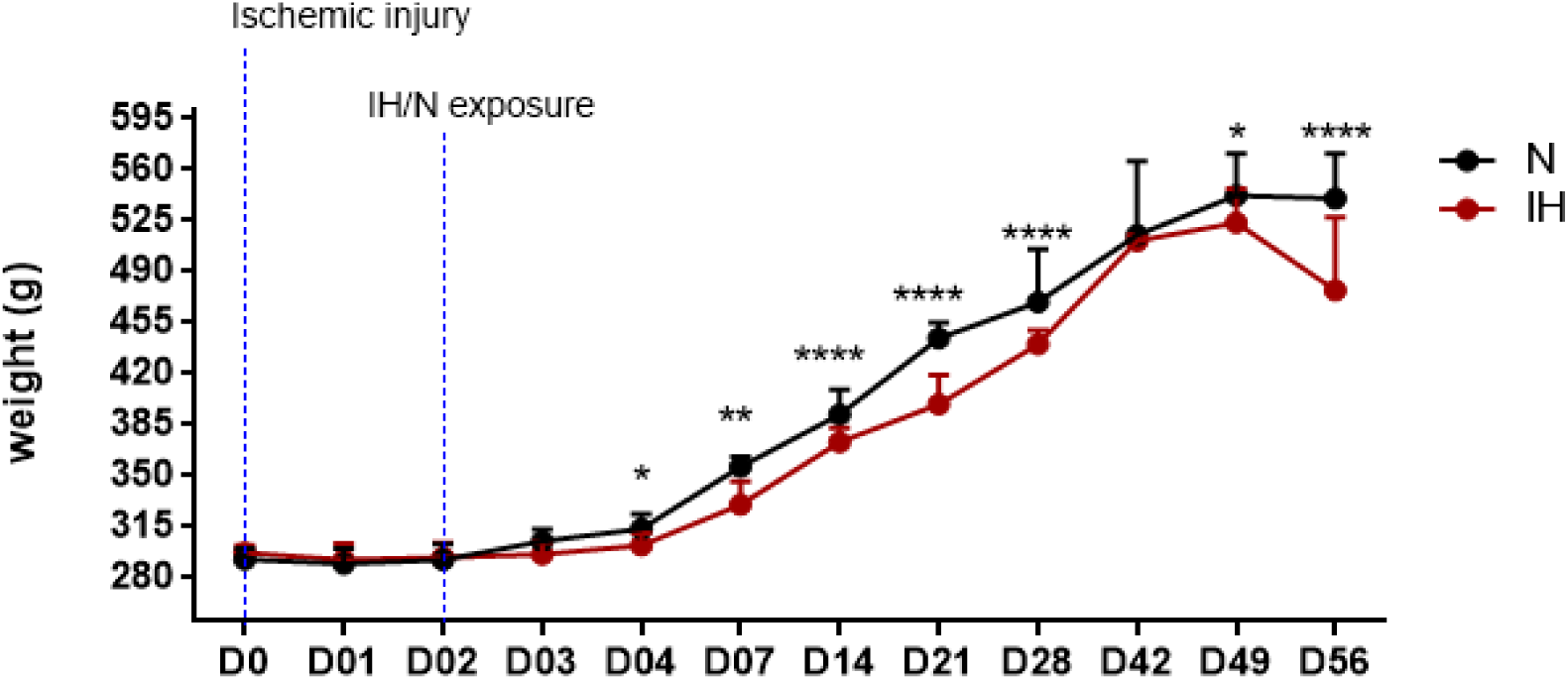
Body weight measurements over 56 days after ischemic injury and IH/N exposure. Body weight (g) was significantly different between IH and N starting at D04 and continued until D49 (2 way-anova). n= 36 N/ 33 IH at D0, n= 36 N/ 32 IH at D1, n= 35 N/ 29 IH at D2, n= 36 N/ 33 IH at D3, n= 28 N/ 33 IH at D04, n= 21 N/ 21(IH at D07, n= 21 N/ 20 IH at D14, n= 8 N/ 9 IH at D21, n= 14 N/ 16 IH at D28, n= 5 N/ 5 IH at D042, n= 5 N/ 5 IH at D049, n= 7 N/ 7 IH at D56. Data are presented with median and interquartile range. * indicates the effect of IH after ischemic injury. * p=0.01, ** p= 0.001, **** p<0.0001.

**Supplementary figure S2.**
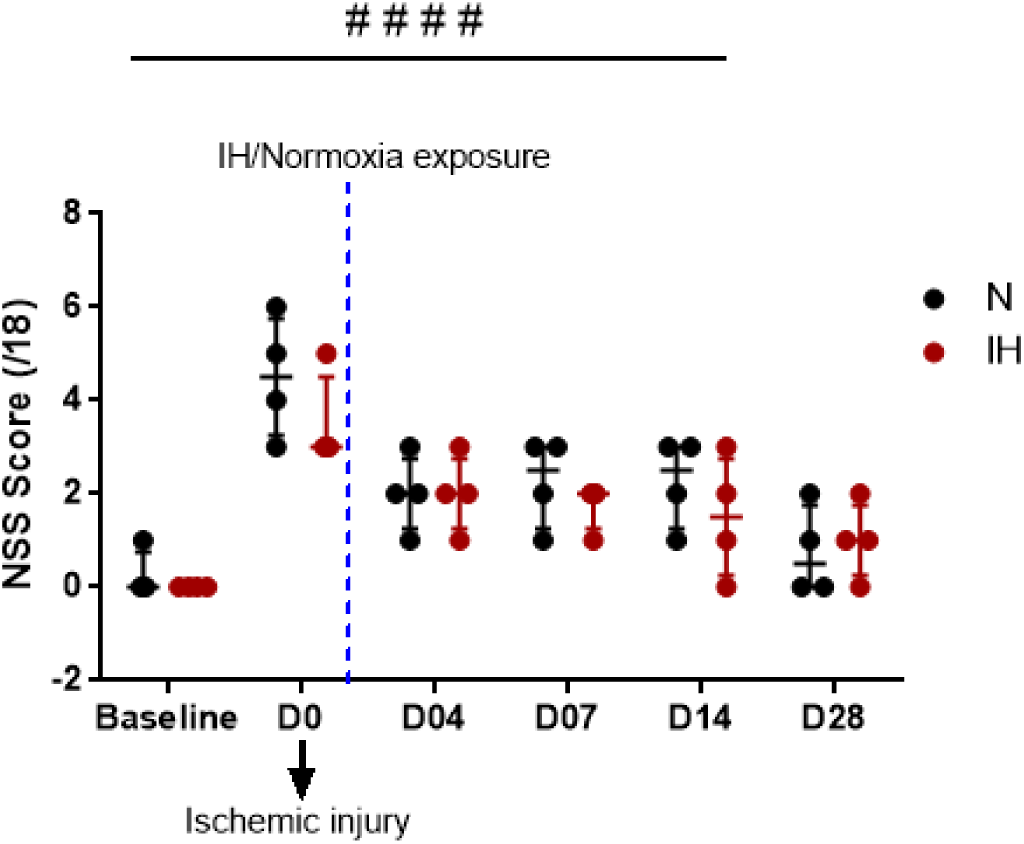
Modified NSS score. NSS score, adopted from [26] performed 2 hours after the surgery to ensure the succession of the ischemic injury. n=4 in each group. Represented data are medians with interquartile range. # indicates the time effect. #### p<0.0001, two-way Anova.

**Supplementary figure S3.**
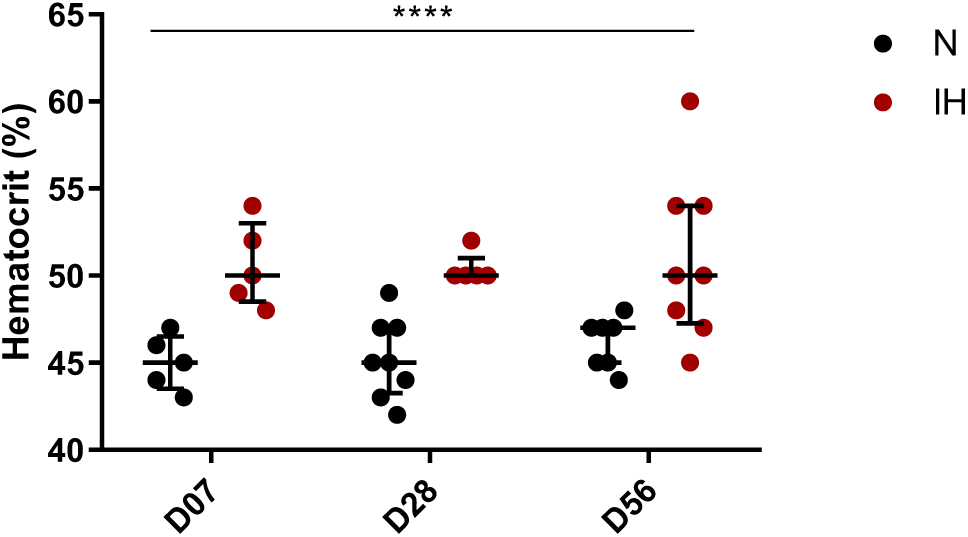
Hematocrit measurements. Hematocrit level was measured at D07 (n= 5N/5 IH), D28 (n= 8 N/6 IH), and D56 (n= 7 N/ 7 IH). Represented data are medians with interquartile range. **** p<0.0001 for the effect of IH, two-way Anova.

**Supplementary table S1:** Overview of MRI timepoints and euthanasia timeline in the 65 experimental rats. MRI protocol 1 : T2 and ADC only; MRI protocol 2 : T2, ADC, R, BVF, O_2_, signal enhancement.

| Rat group and number | Interventions |  |  |  |  |
| --- | --- | --- | --- | --- | --- |
|  | D04 | D07 | D14 | D28 | D56 |
| N (n=5)<br>IH (n=5) | MRI Protocol 2<br>+ euthanasia |  |  |  |  |
| N (n=5)<br>IH (n=4) |  | MRI Protocol 2<br>+ euthanasia |  |  |  |
| N (n=8)<br>IH (n=8) |  | MRI Protocol 2 | MRI Protocol 2<br>+ euthanasia |  |  |
| N (n=8)<br>IH (n=8) |  |  | MRI Protocol 1 | MRI Protocol 2<br>+ euthanasia |  |
| N (n=7)<br>IH (n=7) |  |  |  | MRI Protocol 1 | MRI Protocol 2<br>+ euthanasia |

**Supplementary table 2.** Antibodies used for immunohistological staining.

| Name | Company | Reference | Dilution | Host species | Secondary Antibody | Company | Reference | Dilution |
| --- | --- | --- | --- | --- | --- | --- | --- | --- |
| Collagen 4<br>(**ColIV**) | Abcam | Ab6586 | 1/500 | Rabbit | A568 goat<br>anti-rabbit | Invitrogen | A11036 | 1/500 |
| Neuronal<br>Nuclei<br>(**NeuN**) | Merck<br>millipore | MAB377 | 1/1000 | Mouse | A488<br>donkey<br>anti-mouse | Invitrogen | A21202 | 1/200 |
| Zonula<br>Occludens 1<br>(**ZO-1**) | Thermo<br>Fisher | 40-2200 | 1/200 | Rabbit | A568 goat<br>anti-rabbit | Invitrogen | A11036 | 1/500 |
| Glial fibrillary<br>acidic protein<br>(**GFAP**) | Agilent | Z0334 | 1/1000 | Rabbit | A568 goat<br>anti-rabbit | Invitrogen | A11036 | 1/1000 |

**Supplementary table S3.**
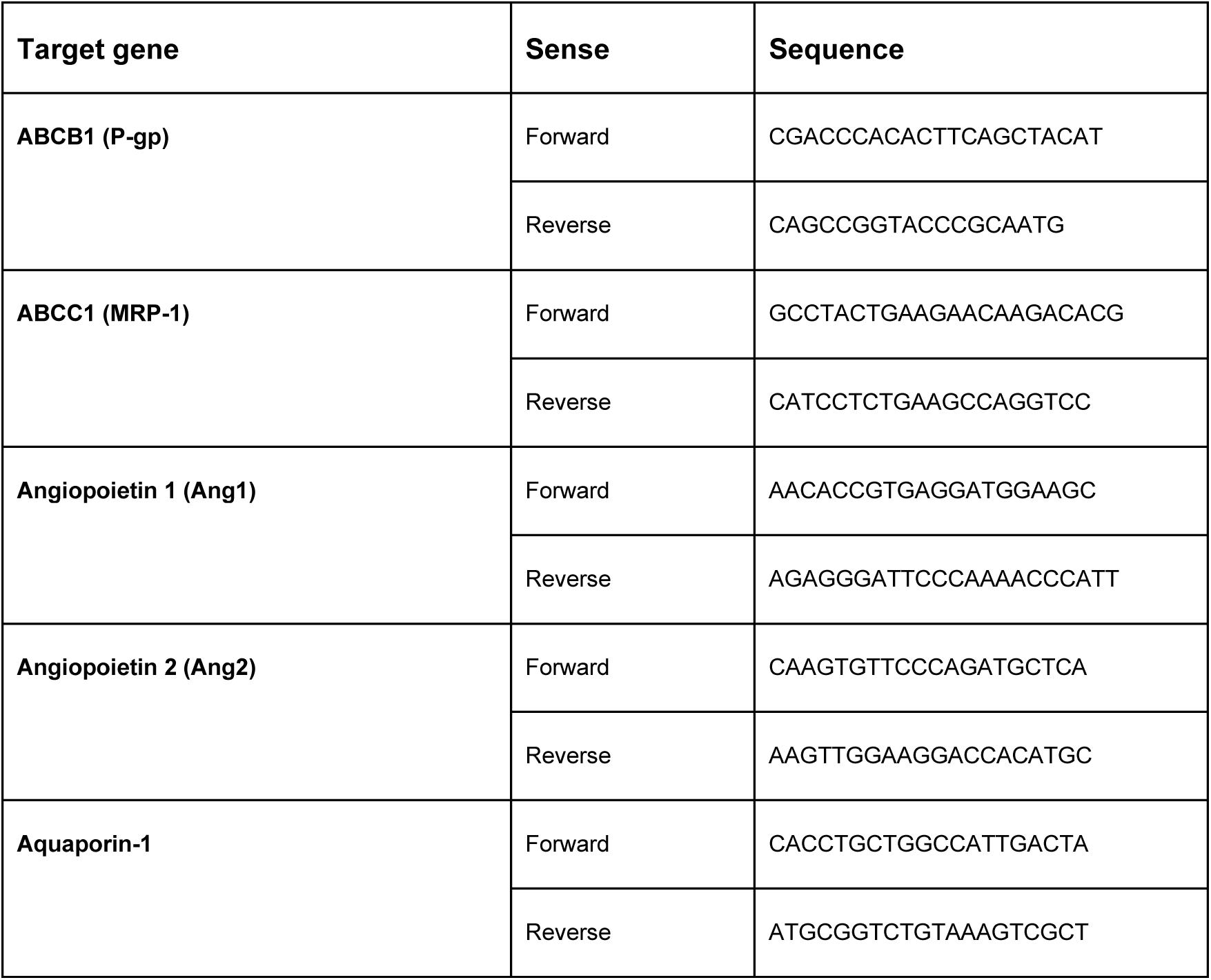
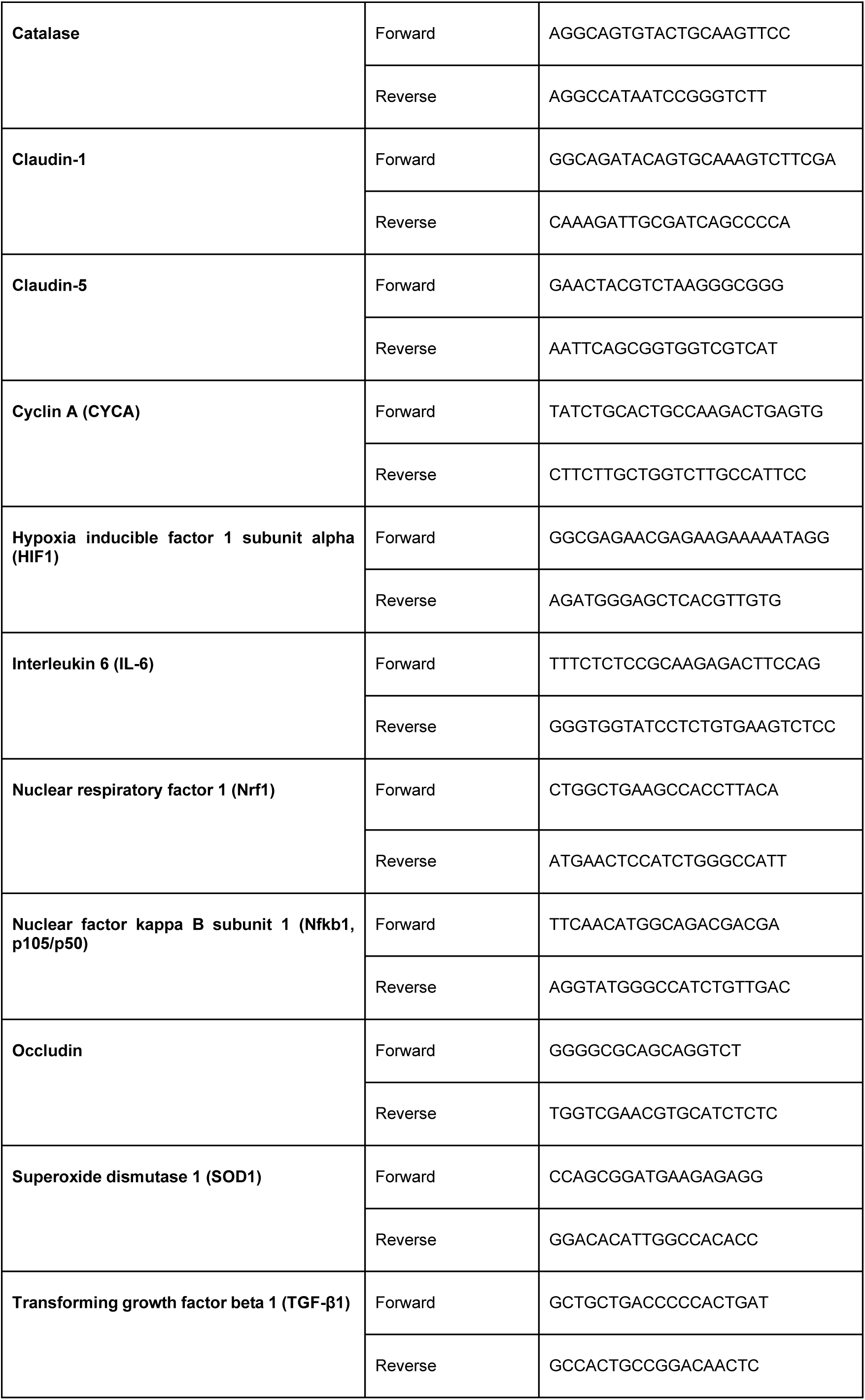
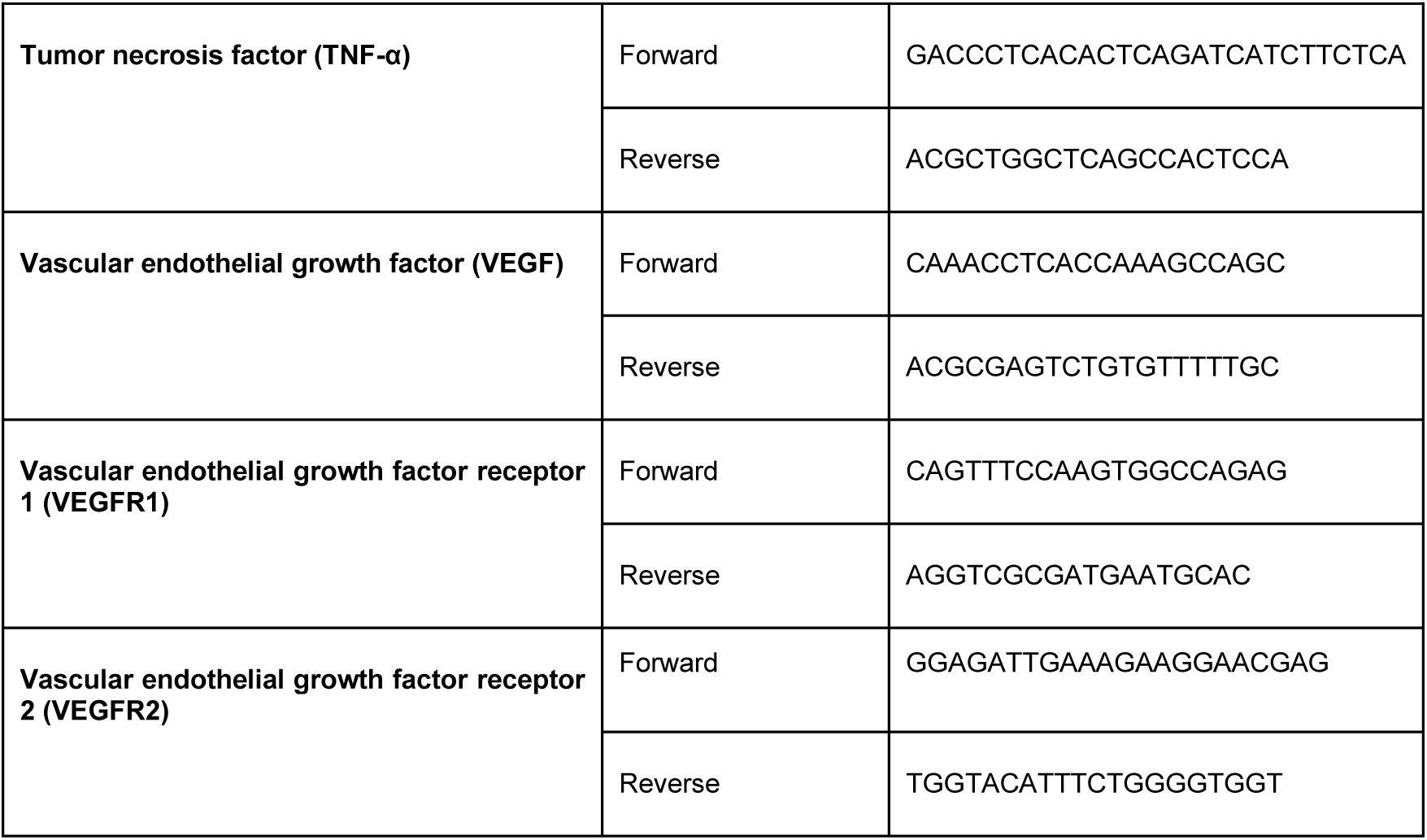
Primer list for qPCR.

**Supplementary table S4.** Reported p-values for the multiple comparisons between consecutive days in IH condition. These post-hoc tests were performed when a significant interaction was found between IH, malonate and time in the three-way ANOVA.

| Gene | VEGF |  | VEGFR1 |  | VEGFR2 |  | Ang2 |  |
| --- | --- | --- | --- | --- | --- | --- | --- | --- |
| Days | D07 | D28 | D07 | D28 | D07 | D28 | D07 | D28 |
| D04 | **0.025** | 0.266 | **0.004** | 0.904 | 0.113 | 0.078 | 0.99 | **0.019** |
| D07 | - | **0.0001** | - | **0.0001** | - | **0.0001** | - | 0.310 |

| Protein | ZO-1 |  | Claudin 1 | Aquaporin 1 |
| --- | --- | --- | --- | --- |
| Days | D07 | D28 | D07 | D07 |
| D04 | 0.916 | 0.96 | 0.99 | 0.99 |
| D07 | - | 1.00 | - | - |

